# Membrane stiffness is a key determinant of *E coli* MscS channel mechanosensitivity

**DOI:** 10.1101/790501

**Authors:** Feng Xue, Charles D. Cox, Navid Bavi, Paul R Rohde, Yoshitaka Nakayama, Boris Martinac

## Abstract

Prokaryotic mechanosensitive (MS) channels have an intimate relationship with membrane lipids. Membrane lipids may influence channel activity by directly interacting with bacterial MS channels or by influencing the global properties of the membrane such as area stretch and bending moduli. Previous work has implicated membrane stiffness as a key determinant of the mechanosensitivity of *E. coli* (*Ec)*MscS. Here we systematically tested this hypothesis using patch fluorometry of azolectin liposomes doped with lipids of increasing area stretch moduli. Increasing DOPE content of azolectin liposomes causes a rightward shift in the tension response curve of *Ec*MscS. These rightward shifts are further magnified by the addition of stiffer forms of PE such as the branched chain lipid DPhPE and the fully saturated lipid DSPE. Furthermore, a comparison of the branched chain lipid DPhPC to the stiffer DPhPE showed a rightward shift in the tension response curve in the presence of the stiffer DPhPE. We show that these changes are not due to changes in membrane bending rigidity as the tension threshold of *Ec*MscS in membranes doped with PC18:1 and PC18:3 are the same, despite a two-fold difference in their bending rigidity. We also show that after prolonged pressure application sudden removal of force in softer membranes causes a rebound reactivation of *Ec*MscS and we discuss the relevance of this phenomenon to bacterial osmoregulation. Collectively, our data demonstrate that membrane stiffness is a key determinant of the mechanosensitivity of *Ec*MscS.

## Introduction

The *E. coli* mechanosensitive (MS) ion channel of small conductance (*Ec*MscS) is a prototypical membrane tension sensor which plays a pivotal role in osmoregulation^*1, 2*^. This channel is the canonical member of a diverse family of MS channels that spans prokaryotic and eukaryotic cell-walled organisms^*3-5*^. Purification and reconstitution of *Ec*MscS, and many of its homologues, into lipid bilayers show that it gates according to the force-from-lipid principle^*6-8*^. This means the channel is inherently mechanosensitive and directly senses membrane forces that result in a conformational change culminating in the channel opening. As a result, it is clear that membrane lipids are a key driver of *Ec*MscS activity. Recent evidence suggests that eukaryotic MS channels also employ force-from-lipids gating^*9-12*^. Therefore, the basic biophysical principles that govern the gating of prokaryotic channels may in turn provide insight into the gating of eukaryotic MS channels^*13, 14*^.

Liposomal reconstitution has for many years been successfully used not only to document the inherent mechanosensitivity of both prokaryotic and eukaryotic ion channels but also to probe the influence of individual lipids on MS channel function^*15-17*^. Lipids can influence integral membrane proteins such as MS channels in one of two ways^*18*^. Firstly, the lipid may directly interact with the protein and modify function. The second is that lipids may indirectly affect function via global effects on the mechanical properties of the bilayer. For example, liposomal reconstitution has shown that the bacterial channel MscL is sensitive to bilayer thickness^*19-21*^, a global property of the bilayer and that the *Mycobacterium tuberculosis* MscL homologue specifically interacts with phosphatidylinositol lipids^*17*^. In comparison bilayer thickness has little effect on *Ec*MscS^*21*^. We are only beginning to understand the structural basis of how lipids directly interact with *Ec*MscS^*22, 23*^ but previous work suggests this channel may be affected by global changes in bilayer stiffness. This is particularly evident when *Ec*MscS is reconstituted into bilayers containing increasing levels of cholesterol, a lipid that increases the area expansion modulus of bilayers^*21*^. Increasing levels of cholesterol cause the pressure threshold of *Ec*MscS to increase. However, many of these studies looking at the sensitivity of *Ec*MscS use applied hydrostatic pressure as a surrogate for in plane membrane tension, the parameter that has been shown to correlate most closely with channel activation^*24-26*^. Furthermore, Piezo1 a eukaryotic mechanosensitive channel that also senses membrane forces has recently been shown to be sensitive to membrane stiffness^*27-29*^.

Here we investigated how the gating kinetics and tension sensitivity of *Ec*MscS were affected by adding defined amounts of lipids with different area expansion moduli to azolectin membranes using patch fluorometry, a technique that combines confocal microscopy and patch clamp electrophysiology^*21, 26*^. A small amount of a fluorescent lipid such as rhodamine PE (<0.1% w/w), was mixed with the desired lipids to accurately follow membrane deformation during patch clamping. By measuring the radius of curvature, the in-plane membrane tension could then be calculated using Laplace’s law^*30*^.

Initially, we aimed to measure the tension sensitivity of purified *Ec*MscS in lipid bilayers composed of dioleoylphosphatidylethanolamine (DOPE) and dioleoylphosphatidylcholine (DOPC) alone. However, none of the combinations of DOPE/DOPC were amenable to the pressure protocol required to accurately measure membrane tension. As a result, we employed azolectin liposomes mixed with lipids of different area expansion modulus. Given that all stiffer lipids caused a significant rightward shift of the *Ec*MscS tension response curve, and increased the channel tension threshold, our data clearly shows that membrane stiffness is a key determinant of the mechanosensitivity of *Ec*MscS.

## Methods

### Lipids

This study utilized soybean azolectin from Sigma-Aldrich (P5638). 1,2-dioleoyl-sn-glycero-3-phosphocholine (DOPC), 1,2-Dioleoyl-sn-glycero-3-phosphoethanolamine (DOPE), 1,2-diphytanoyl-sn-glycero-3-phosphocholine (DPhPC), 1,2-diphytanoyl-sn-glycero-3-phosphoethanolamine (DPhPE), 1,2-distearoyl-sn-glycero-3-phosphocholine (DSPC), 1,2-Distearoyl-sn-glycero-3-phosphoethanolamine (DSPE), 1,2-dipalmitoleoyl-sn-glycero-3-phosphoethanolamine (PE 16:1) and 1,2-dioleoyl-sn-glycero-3-phosphoethanolamine-N-(lissamine rhodamine B sulfonyl) were purchased from Avanti.

### *E. coli* MscS protein purification and reconstitution

*E. coli* MscS was purified using a 6xHis-tag according to previously published protocols^*31*^. Prior to reconstitution the 6xHis-tag was cleaved with thrombin and *Ec*MscS was reconstituted into liposomes with different lipid components using the dehydration/rehydration (D/R) reconstitution method. Azolectin was dissolved in chloroform and mixed with the respective lipids of interest. Fluorescent rhodamine-PE is then added at 0.1% before the lipid mixture is dried under nitrogen. The lipid film was then suspended in D/R buffer (200mM KCl, 5mM HEPES, pH adjusted to 7.2 using KOH) and vortexed followed by water bath sonication (6 L 120 W pulse swept power) for 15 minutes. Then 1:50 (w:w) MscS protein was added into the lipid mixture and incubated for 1 hour with agitation, followed by the addition of 300 mg of Biobeads (SM-2, BioRad). The Biobeads were mixed for three hours at room temperature. Finally, the mixture was ultracentrifuged at 40,000 RPM in a Beckman Type 50.2 Ti rotor for 30 mins and the lipid mixture was vacuum desiccated overnight. The protein reconstituted liposomes were rehydrated in D/R buffer overnight before use.

### Electrophysiology

Liposomes were incubated in patch buffer containing: 200mM KCl, 40mM MgCl2, 5mM HEPES adjusted to pH 7.2 using KOH, for one hour until unilamellar blisters formed on their surface. The patch pipette solution and bath solution were symmetric in all recordings containing, in mM: 200 KCl, 40 MgCl2 and 5 HEPES-KOH, pH 7.2. The single channel currents were amplified using an Axopatch 200B amplifier (Molecular Devices). The *E. coli* MscS currents were filtered at 2 kHz and sampled at 5 kHz with a Digidata 1440A using pClamp 10 software. Negative hydrostatic pressure was applied in 1s increments by -10 mmHg via a high-speed pressure clamp (ALA Sciences) up to a maximum pressure of -100 mmHg.

### Patch fluorometry

Wild-type *Ec*MscS channels were added to liposomes at a protein:lipid ratio of 1:50 (w/w) and recorded by imaging the tip of the patch pipette using a confocal microscope (LSM 700; Carl Zeiss) housed within a Faraday cage and equipped with a water immersion objective lens (×63, NA1.15). The excised liposome patches that consisted of 99.9% lipids of interest and 0.1% lissamine rhodamine-phosphatidylethanolamine (PE) (w/w) were excited with a 555 nm laser. Fluorescence images of the deformed membranes were acquired and analyzed with ZEN software (Carl Zeiss GmbH). To improve visualization of liposome patches even further the pipette tip was bent approximately 30° with a microforge (MF-900; Narishige, Tokyo, Japan) to make it parallel to the bottom face of the recording chamber. The diameter of the patch dome at each step of pressure was measured using the ZEN software. Tension is calculated using Laplace’s law as previously described^19, 27^.

## Results and discussion

### Sensitivity of *E. coli* MscS in pure DOPC/DOPE bilayers

We first attempted to accurately measure the tension sensitivity of *E. coli* MscS (*Ec*MscS) in bilayers comprised of DOPC and DOPE. This is due to the fact that previous work shows that membranes composed of PE are stiffer than those of PC^*32*^ and that the rigidity of PC membranes can be increased by the sequential addition of PE^*33*^. However, we found that the long stepwise pressure (1 s × 10 steps) application via a high-speed pressure clamp necessary to accurately measure the curvature of the membrane was not compatible with pure DOPC/DOPE bilayers (Fig. 1A-C).

**Figure 1.**
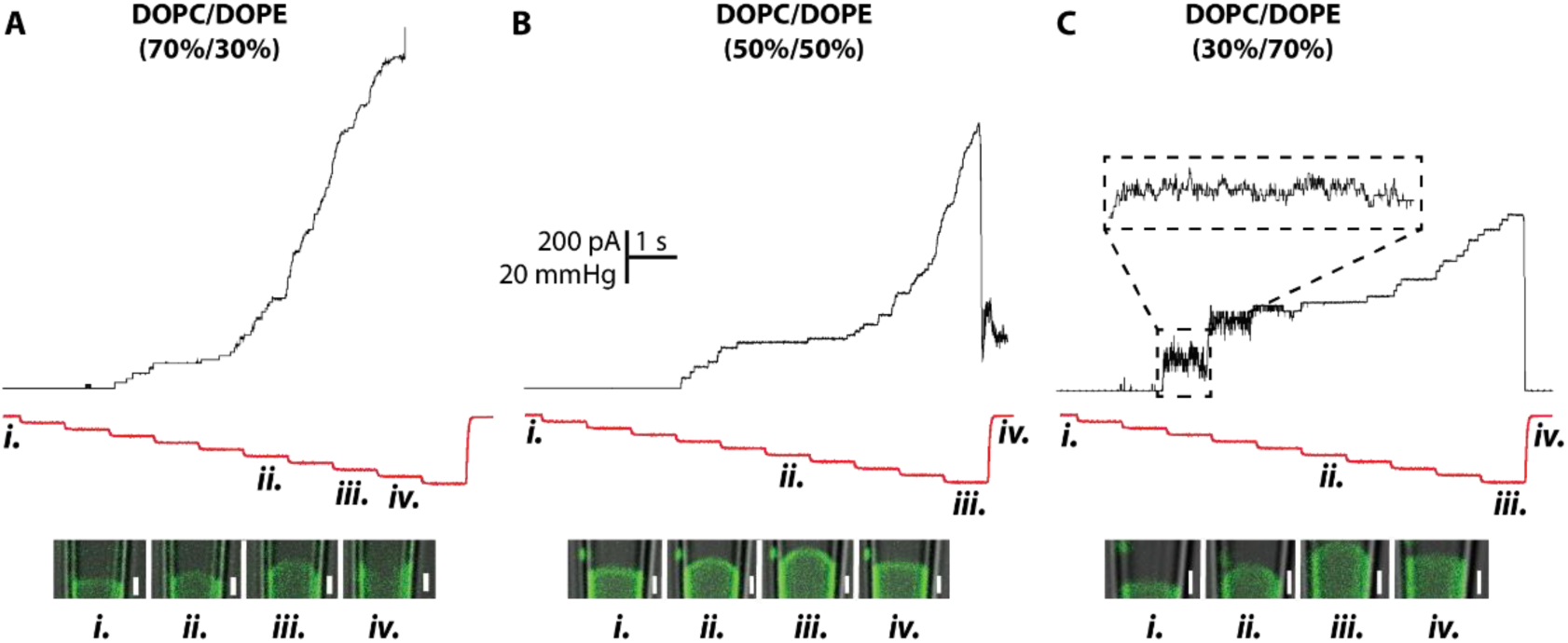
EcMscS gating in pure DOPC/DOPE liposomal membranes. (A) Representative patch clamp recording of EcMscS reconstituted in DOPC/DOPE (70%/30%). (B) Representative patch clamp recordings of EcMscS reconstituted in DOPC/DOPE (50%/50%). (C) Representative patch clamp recording of EcMscS reconstituted in DOPC/DOPE (30%/70%), inset shows increased gating events in this lipid mixture. Black trace represents current and negative pressure steps applied via high speed pressure clamp are shown in red. Replicates and statistics associated with DOPC/DOPE liposomes are shown in Table 1.

The details of the replicates from DOPC/DOPE bilayers are shown in Table.1. In the group of *Ec*MscS reconstituted into DOPC/DOPE (70%/30%) liposomes, all channels in the patch could be activated but this only occurred in 1 out of 16 patches with 12/16 not surviving the full step protocol (Fig.1A, Table.1). When the concentration of DOPE was increased the ‘survival rate’ of membrane patches under the pressure protocol [1s steps in increments of 10mmHg with a total of 10 steps] was higher. However, in both DOPC/DOPE (50%/50%) and DOPC/DOPE (30%/70%), the saturation point (peak current) of all *Ec*MscS in the patch could not be reached prior to the membrane reaching its physical limit. Thus, we could not reach peak current of *Ec*MscS and consequently could not accurately measure the tension sensitivity of *Ec*MscS in these membranes. However, we did notice that the gating of *Ec*MscS was more ‘flickery’ in 70%/30% as previously reported (Fig.1C). We also attempted patch clamp experiments with *Ec*MscS reconstituted into DOPC (100%) but in all attempts membranes broke before pressure was applied (0/20). In agreement with earlier liposome studies we were unable to form unilamellar blisters in DOPE only (100%).

**Table 1.** Statistics associated with patch fluorometry experiments using pure DOPC and DOPE liposomal membranes. The table documents the total number of trials, the number of patches where EcMscS channel activity did not reach a plateau despite reaching -100 mmHg, the number of patches that ruptured before all channels were activated and the number of patches where the maximal number of EcMscS channels in the patch were activated.

| Lipid Composition | Peak current not reached | Rupture before peak current | Peak current reached | Total trials |
| --- | --- | --- | --- | --- |
| DOPC 100% | N/A | 20 | N/A | 20 |
| DOPC/DOPE 70/30 | 3 | 12 | 1 | 16 |
| DOPC/DOPE 50/50 | 7 | 5 | 0 | 12 |
| DOPC/DOPE 70/30 | 11 | 2 | 0 | 13 |

### The effect of DOPE on *E. coli* MscS threshold

In order to overcome the issues associated with our pressure protocol and the fragility of DOPC/DOPE membranes we decided to use azolectin liposomes doped with lipids of interest and consequently tested the tension sensitivity of *Ec*MscS. We chose azolectin as it has been widely used to reconstitute eukaryotic and prokaryotic ion channels^*11, 21, 34, 35*^. Azolectin is a lipid mixture primarily composed of PC and as previously mentioned the sequential addition of PE stiffens PC membranes^*33*^. Given that PE is the major constituent of *E. coli* membranes and DOPE is inherently stiffer than DOPC^*32*^ we first looked at the effect of increasing amounts of DOPE on *Ec*MscS gating in azolectin liposomes. Firstly, our results show that when the concentration of DOPE is above 30% in the Azolectin (Azo)/DOPE mixture, the gating of *Ec*MscS became ‘flickery’. Flickering was observed in Azo/DOPE (40%/60%) and Azo/DOPE (70%/30%) groups before the peak current was reached (Fig. 2C & 3A). This increase in the number of gating events has been documented and quantified previously^*15*^.

**Figure 2.**
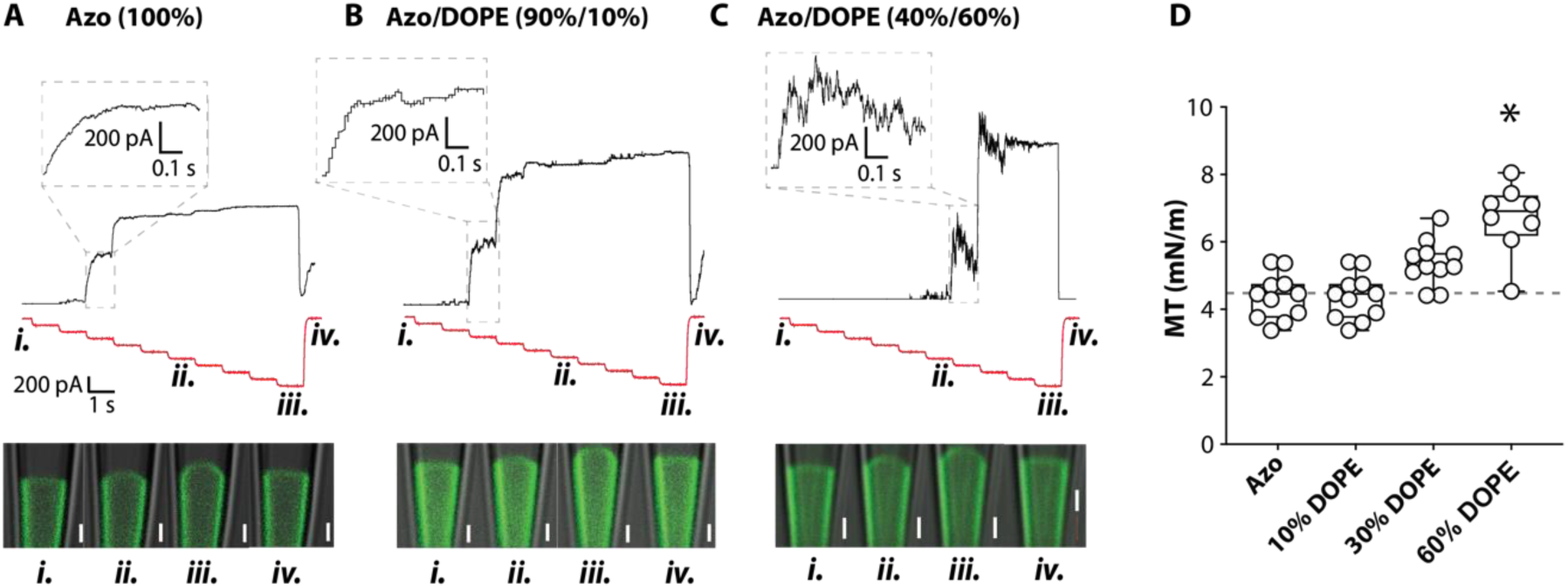
The effects of increasing concentrations of DOPE on the tension sensitivity of EcMscS. **(A)** Patch clamp recordings of EcMscS with corresponding patch fluorometry images documenting Azolectin (Azo) bilayer deformation (n=8). **(B)** Patch clamp recordings of EcMscS with corresponding patch fluorometry images documenting Azo/DOPE(90%/10%) bilayer deformation (n=11). (C) Patch clamp recordings of EcMscS with corresponding patch fluorometry images documenting Azo/DOPE(40%/60%) bilayer deformation (n=8). **(D)** Activation curves of EcMscS in liposome membranes containing different amounts of DOPE. The activation curves of EcMscS were shifted towards right with increasing the amount of DOPE. Box and whiskers plot illustrating the midpoint tension (MT) of EcMscS reconstituted in Azolectin, Azo/DOPE(90%/10%), Azo/DOPE(70%/30%) and Azo/DOPE(60%/40%). * denotes statistically significant difference from azolectin alone using Kruskal-Wallis with Dunn’s post hoc test, p < 0.05. Inset white bar on images represents 2 µm.

**Figure 3.**
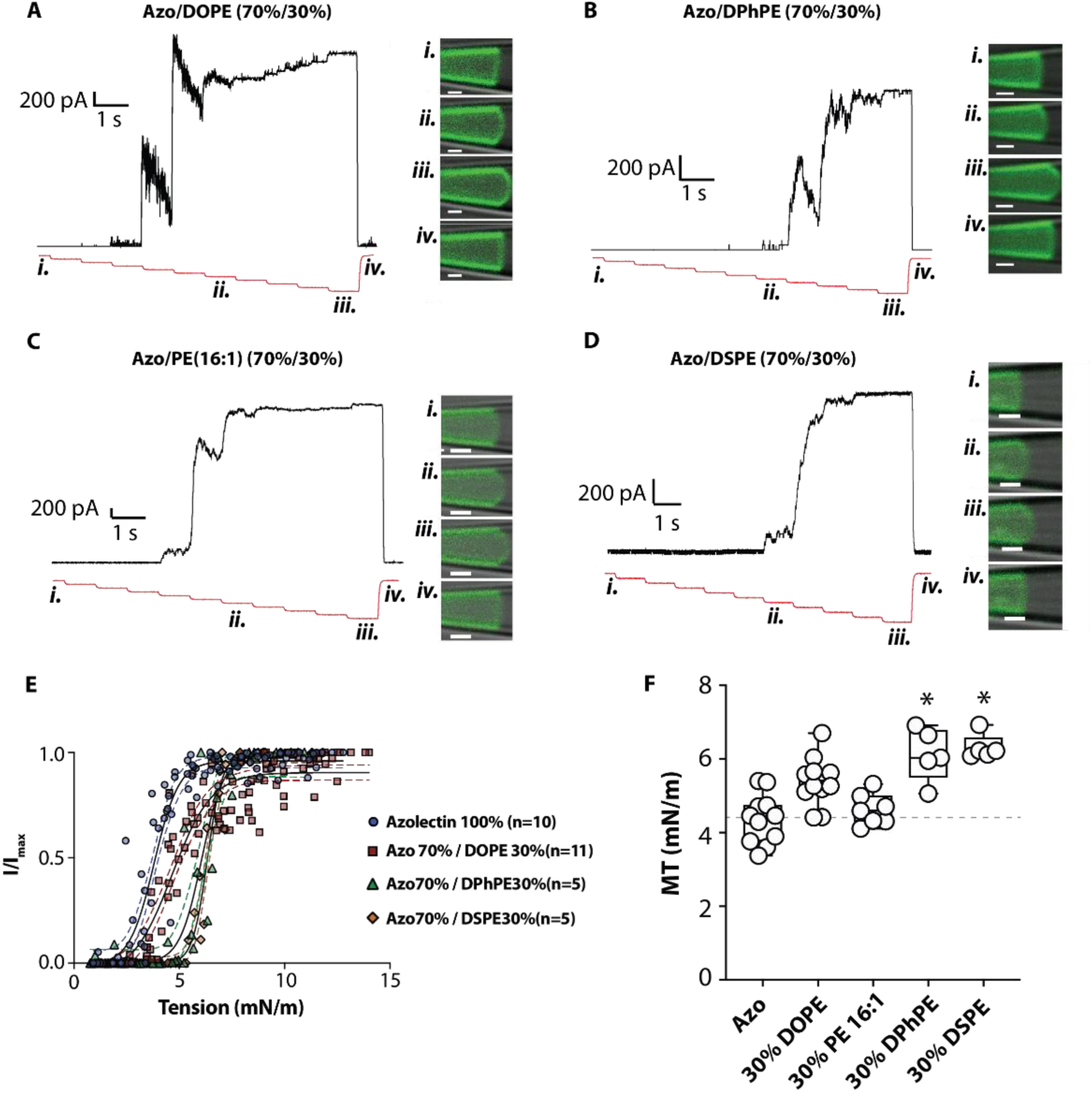
The effect of DOPE, DPhPE, PE(16:1), and DSPE on the tension sensitivity of EcMscS. **(A)** Patch clamp recordings of EcMscS with corresponding patch fluorometry images documenting Azolectin/DOPE (PE18:1) (70%/30%) bilayer deformation (n=11). **(B)** Patch clamp recordings of EcMscS with corresponding patch fluorometry images documenting Azolectin/DPhPE (70%/30%) bilayer deformation (n=5). **(C)** Patch clamp recordings of EcMscS with corresponding patch fluorometry images documenting Azolectin/PE (PE16:1) (70%/30%) bilayer deformation (n=11). **(D)** Patch clamp recordings of EcMscS with corresponding patch fluorometry images documenting Azolectin/DSPE (PE18:0) (70%/30%) bilayer deformation (n=5). **(E)** Tension response curves of EcMscS in all groups compared to azolectin alone shown in blue. Coloured lines represent the 95% confidence intervals for the Boltzmann fits shown in black. **(F)** Box and whiskers plot showing the midpoint tension threshold (MT) of EcMscS in each lipid group. * denotes statistically significant difference from azolectin alone using Kruskal-Wallis with Dunn’s post hoc test p < 0.05. Inset white bar on images represents 2 µm.

From the activation curve of *Ec*MscS, when the concentration of DOPE in the Azo/DOPE mixture increased, the midpoint tension also increased (Fig.2A-D). The midpoint tension of *Ec*MscS reconstituted into Azolectin (100%) was 4.4 ± 0.2 mN/m while the threshold in Azo/DOPE (90%/10%), Azo/DOPE (70%/30%), Azo/DOPE (40%/60%) was 4.8 ± 0.4 mN/m, 5.3 ± 0.2 mN/m and 6.7 ± 0.4 mN/m, respectively. These values are similar to previous estimates for the midpoint tension of *Ec*MscS^*21, 26*^. These results were not affected by the channel number, because both the average channel number in Azolectin (100%), in which the channels required the least tension to open, and Azo/DOPE(40%/60%), in which the channels required the most tension to open, were similar (Table 2). Importantly, in all our data sets there is no correlation between the number of channels incorporated and the midpoint tension threshold (Table 2).

**Table 2.** Channel number, slope and midpoint threshold in all azolectin lipid compositions tested in this study. Data represents mean ± SEM.

| Lipid Composition | N | Channel No. | Slope (-mN/m) | MT (mN/m) |
| --- | --- | --- | --- | --- |
| Azo 100% | 11 | 46 ± 7 | 0.59 ± 0.08 | 4.4 ± 0.2 |
| Azo/DOPE 90/10 | 11 | 89 ± 17 | 0.58 ± 0.09 | 4.8 ± 0.4 |
| Azo/DOPE 70/30 | 11 | 42 ± 6 | 0.79 ± 0.10 | 5.3 ± 0.2 |
| Azo/DOPE 40/60 | 8 | 46 ± 8 | 0.70 ± 0.11 | 6.7 ± 0.4 |
| Azo/PE 16:1 70/30 | 7 | 83 ± 31 | 0.42 ± 0.04 | 4.6 ± 0.5 |
| Azo/DPhPE 40/60 | 5 | 44 ± 19 | 0.69 ± 0.17 | 6.1 ± 0.3 |
| Azo/DSPE 70/30 | 5 | 59 ± 16 | 0.36 ± 0.04 | 6.3 ± 0.2 |
| Azo/DOPC 70/30 | 9 | 17 ± 5 | 0.67 ± 0.13 | 2.4 ± 0.2 |
| Azo/PC18:3 70/30 | 5 | 32 ± 12 | 0.48 ± 0.16 | 2.6 ± 0.2 |
| Azo/DSPC 70/30 | 10 | 0 ± 0 | ND | ND |
| Azo/DPhPC 70/30 | 6 | 38 ± 9 | 0.69 ± 0.17 | 4.8 ± 0.4 |

### The effect of stiffer forms of PE on *E. coli* MscS tension sensitivity

In order to further interrogate the impact of PE on *Ec*MscS tension sensitivity we made use of two forms of PE which are stiffer than DOPE namely DPhPE and DSPE (18:0) (Fig. 3B and D). We found that adding 30% of DPhPE and DSPE to azolectin liposomes caused a significant rightward shift in the tension response curve of *Ec*MscS (Fig. 3E, Table 2). The midpoint tension for *Ec*MscS in the presence of 30% DPhPE and DSPE were 6.1 ± 0.3 and 6.3 ± 0.2 mN/m, respectively. These rightward shifts were larger in magnitude than measured for DOPE 30% (Fig. 3E, Table 2). DPhPE has a carbon chain length of 16 compared to the c18 of DOPE. In order to ensure that this rightward shift was not due to the different thickness we also tested PE 16:1. Previous work suggests that MscL is sensitive membrane thickness^*20*^ but *Ec*MscS is not^*21*^. As expected PE16:1 did not cause the same significant rightward shift in tension sensitivity as DPhPE (Fig. 3C and E, Table2).

The channel flickering also occurred when *Ec*MscS was reconstituted in Azo/DPhPE (70%/30%) (Fig.3B) but not in Azo/DSPE (70%/30%). This suggests that different membrane parameters drive the tension sensitivity changes and the kinetic changes.

### The effect of increasing PC content on *E. coli* MscS threshold

We also undertook analogous experiments using PC. *E. coli* MscS reconstituted into Azo/DOPC(70%/30%) liposomes had a higher sensitivity than in Azolectin alone. The midpoint threshold of *Ec*MscS in Azo/DOPC (70%/30%) liposomes was 2.4 ± 0.2 mN/m which is lower than the midpoint threshold of *Ec*MscS in Azolectin (Fig. 4A, Table 2).

**Figure 4.**
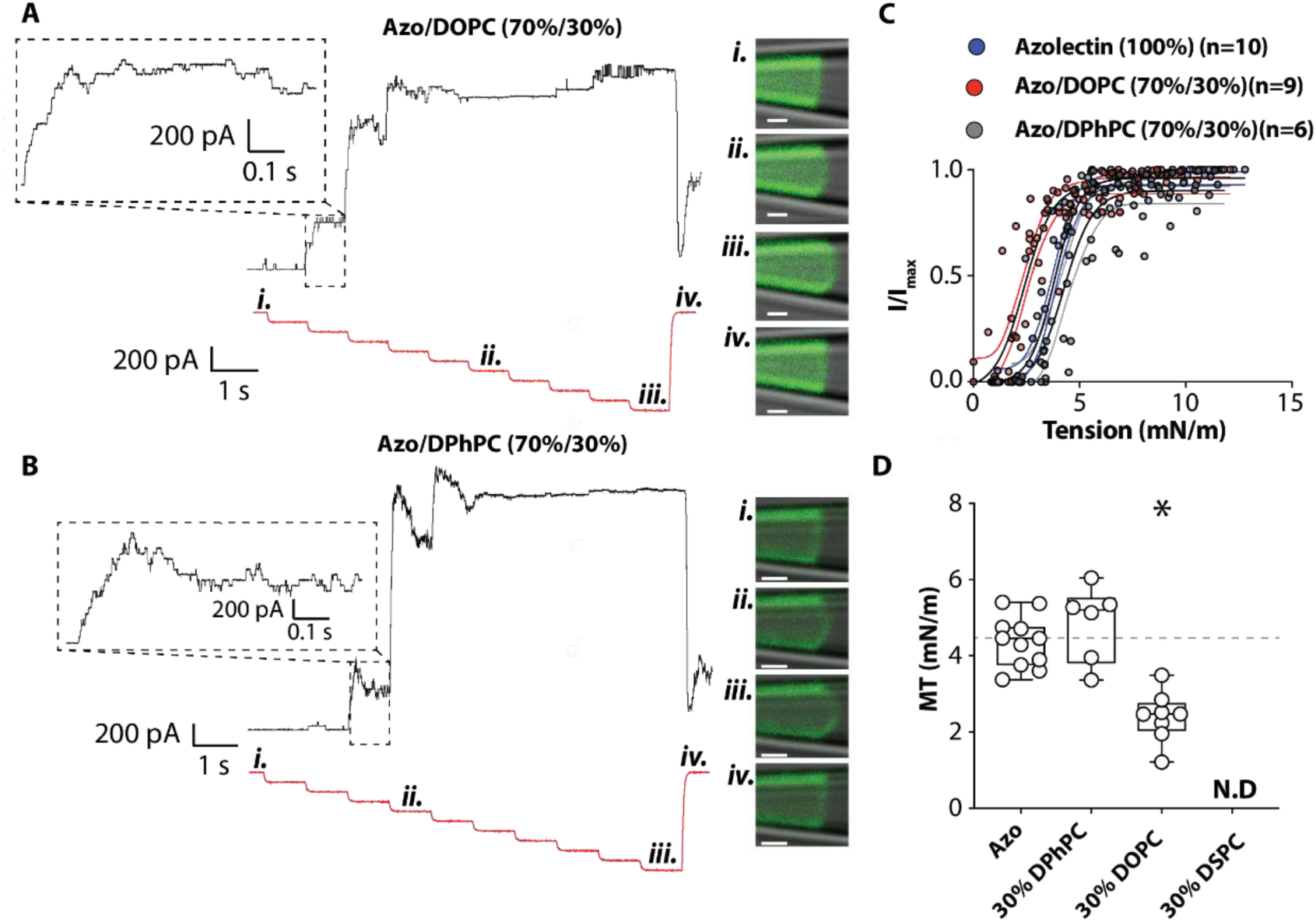
The effect of PC on the tension sensitivity of EcMscS. **(A)** Patch clamp recordings of EcMscS with corresponding patch fluorometry images documenting Azolectin/DOPC (PC18:1) (70%/30%) bilayer deformation (n=8). **(B)** Patch clamp recordings of EcMscS with corresponding patch fluorometry images documenting Azolectin/DPhPC (70%/30%) bilayer deformation (n=5). **(C)** Tension response curves of EcMscS in all groups compared to azolectin alone are shown in blue. Coloured lines represent 95% confidence intervals for the Boltzmann fits shown in black. **(D)** Box and whiskers plot showing the midpoint tension threshold (MT) of EcMscS in Azolectin, Azo/DOPC (70%/30%), Azo/DPhPC (70%/30%) and Azo/DSPC (70%/30%). ND means not determined. * denotes statistically significant difference from azolectin alone using Kruskal-Wallis with Dunn’s post hoc test, p < 0.05. Inset white bar on images represents 2 µm.

The channel number in Azo/DOPC (70%/30%) liposomes was much lower than in almost all other groups tested with one exception. In the case of Azo/DSPC (70%/30%) from 10 patches from three separate reconstitutions we could not identify any *Ec*MscS activity.

We compared the tension sensitivity of *Ec*MscS in 30% DPhPC to 30% DOPC and a clear rightward shift was observed (Fig. 4C). The midpoint tension of 30% DPhPC and DPhPE were 4.8 ± 0.4 mN/m and 6.1 ± 0.3 mN/m, respectively (Fig. 4D). This matches the higher elastic modulus of DPhPE (160 mN/m) when compared to DPhPC (125 mN/m). Thus, all our data is congruent with the idea that stiffer membranes reduce the tension sensitivity of *Ec*MscS.

### *Ec*MscS tension sensitivity does not correlate with bending rigidity

In all of the cases tested the tension sensitivity of *Ec*MscS correlates with both the area expansion moduli of the lipids (higher the area expansion moduli the higher the tension required for channel opening) and the bending rigidity (higher the bending rigidity the higher the tension required for channel opening). In order to see which physical parameter is more important we made use of polyunsaturated acyl chains. In particular we looked at azolectin liposomes doped with PC18:1 and PC18:3. The area expansion moduli of these lipids are almost identical when measured *in vitro* but the bending rigidity of PC18:3 is half that of PC18:1^*36*^. We find that the midpoint tension threshold of *Ec*MscS in Azo/PC18:3(70%/30%) is very similar to that of azo/PC18:1(70%/30%). This provides further evidence that area expansion modulus and not bending rigidity is the key determinant of *Ec*MscS activity.

### Rebound activity of *Ec*MscS after removal of mechanical stimuli

We noted that on many occasions after the removal of pressure there was a rebound reactivation of *Ec*MscS channels. Figures 6A-B show examples of this rebound activity. We sought to quantitate whether this rebound activity was more or less common in certain lipid groups and found that the highest levels of rebound activation were in liposomes composed of PC lipids. As the levels of PE, particularly stiffer PE types such as DSPE (18:0), increased we saw almost no rebound activation of MscS (Fig. 6C).

**Figure 5.**
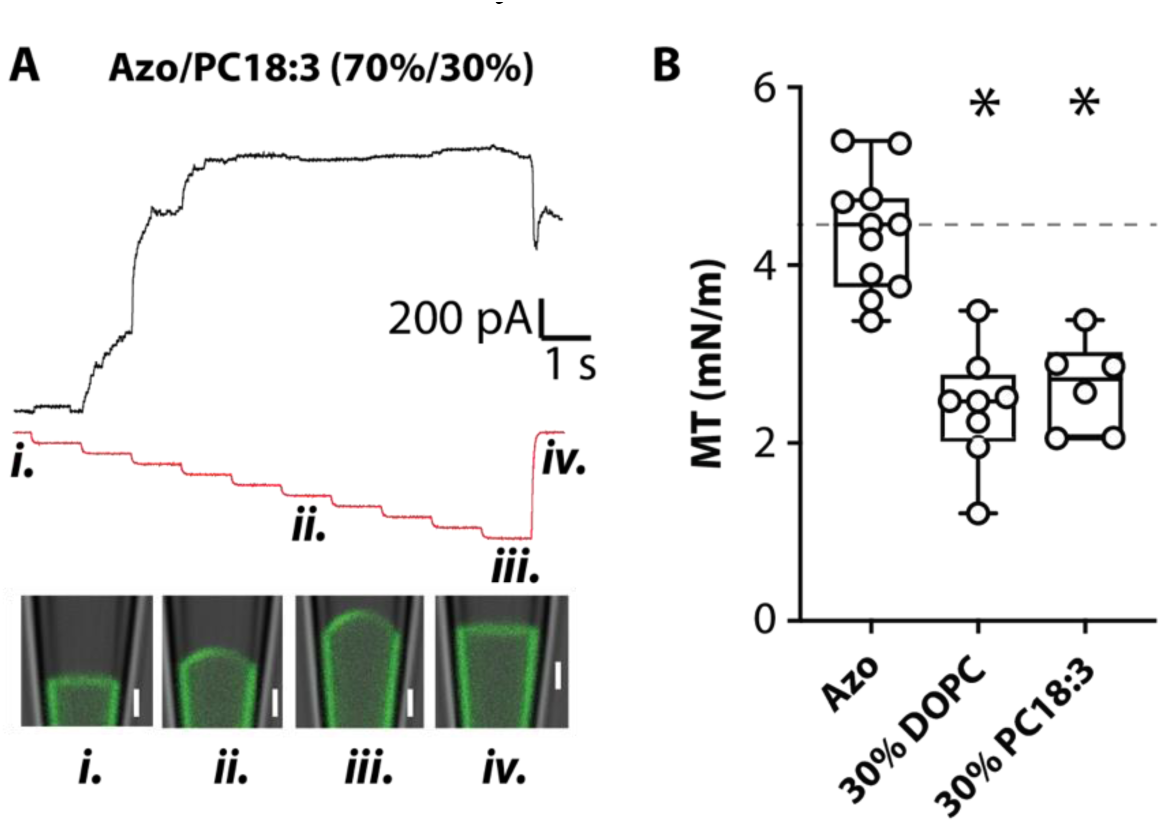
The effect of PC18:3 on the tension sensitivity of EcMscS. **(A)** Patch clamp recordings of EcMscS with corresponding patch fluorometry images documenting Azolectin/PC18:3 (70%/30%) bilayer deformation (n=8). **(B)** Box and whiskers plot showing the midpoint tension threshold (MT) of EcMscS in Azolectin, Azo/DOPC(70%/30%), Azo/PC18:3(70%/30%). * denotes statistically significant difference from azolectin alone using Kruskal-Wallis with Dunn’s post hoc test, p < 0.05. Inset white bar on images represents 2 µm.

**Figure 6.**
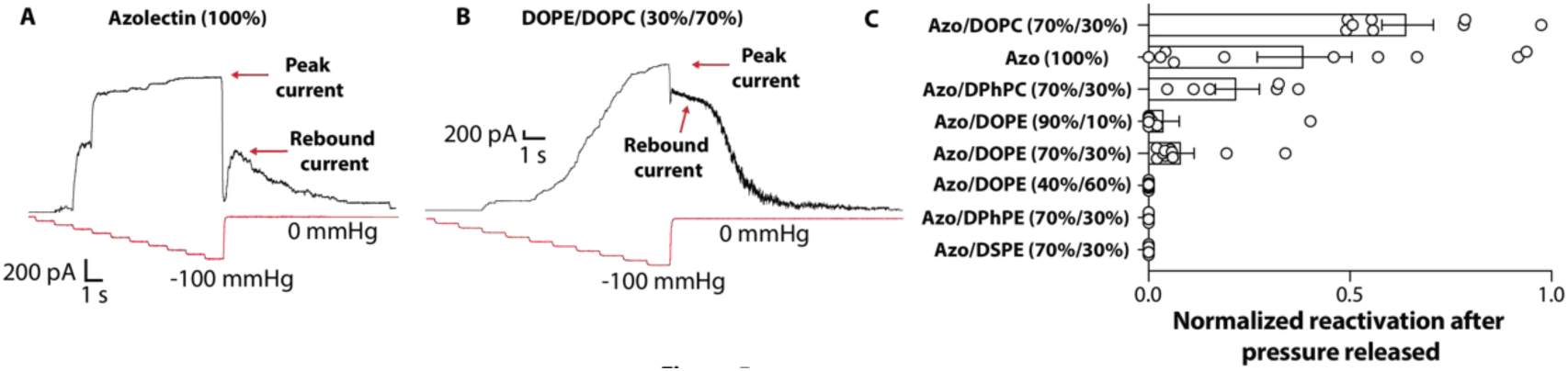
Rebound EcMscS activity after pressure removal. **(A)** Example trace of EcMscS activity reconstituted in Azolectin (100%). Red arrow illustrates the rebound activity where the negative pressure has been released and returns to 0 mmHg. **(B)** The only current trace of EcMscS activity reconstituted in DOPE/DOPC (30%/70%) that progressed through the full negative pressure protocol. Note the exceedingly large rebound current. **(C)** Quantitation of the rebound EcMscS current by normalizing to the peak current. Data represents mean ± SEM (n = 5-11).

This rebound activation seems to correlate well with the midpoint tension thresholds suggesting that the rebound activation increases as the membranes become softer (Fig. 6C). The phenomenon of ‘rebound’ is typical of shock absorbers, which suppress excess force or rapid movement in mechanical systems. Thus, the rebound MscS activity can be described in this way because the lipid bilayer is elastic, but incorporation of protein may introduce viscoelastic properties. The rapid release of pressure/tension (shock) in the liposome bilayer requires the energy accumulated in the now viscoelastic membrane to be dissipated in some way. In the *Ec*MscS case in liposomes this apparently happens through rebound channel activity. The stretching energy stored in the bilayer spring cannot be sufficiently quickly dissipated through the dashpot, which plays a bigger role in soft membranes meaning the softer the membrane the slower becomes the energy dissipation upon sudden release of pressure. Therefore, the rebound effect is larger in softer membranes. This may have implications for *E. coli* osmoregulation and in part explain why their membranes are largely composed of PE. The rapid swelling that occurs in *E. coli* cells as a result of an osmotic downshock is followed by the activation of MS channels that dissipate the generated membrane tension. If the membranes were softer like azolectin or other PC groups, this would result in a rebound activation of the channels and excessive loss of internal solutes. Thus along with the inactivation of the *Ec*MscS channel^*37*^ the PE containing membrane may limit excessive activation, maximizing the chance of survival.

Here, we have avoided the use of charged lipids as these lipids may also be involved in tight protein-lipid interactions with *Ec*MscS residues such as R46^*22*^. Future work should aim to address the contribution of global effects on bilayer mechanics and direct protein-lipid interactions in *Ec*MscS mechanosensitivity.

## Conclusion

Since the major component in Azolectin is PC which forms membranes with a lower Young’s modulus than membranes formed by PE^*32, 33*^, our results suggest that the stiffer membranes make it harder to open *Ec*MscS. Thus, as the PE content becomes larger, we see a rightward shift in the tension response curve of *Ec*MscS channel activity. This pattern is retained when comparing either DOPE with DPhPE or DOPC with DPhPC both of which are much stiffer branched lipids. In fact we also know that DPhPE is much stiffer than DPhPC^*38*^ and again we can see that the 30% DPhPE group has a higher midpoint tension threshold than the 30% DPhPC group. This is highly unlikely to be a chain length effect as our work in addition to previous work has conclusively shown that chain length does not markedly affect *Ec*MscS tension sensitivity, as it does for *Ec*MscL^*21*^. However, when the same amount of azolectin lipid was replaced by lipids which can form softer membranes, the midpoint tension of *E. coli* MscS becomes lower. For example, when the same amount of azolectin was replaced by DOPC, the channel becomes easier to open. This is the same for PC18:1 and 18:3 providing clear evidence that the critical factor is not bending rigidity. In conclusion, our results strongly suggest that membrane stiffness is a key determinant of the mechanosensitivity of *E. coli* MscS channels.

## Author Contributions

The manuscript was written through contributions of all authors. All authors have given approval to the final version of the manuscript.

